# Conceptualization and Feasibility Testing of a Vibro-Acoustic Solution for Tumor Detection

**DOI:** 10.1101/2025.08.23.671745

**Authors:** Mostafa Sayahkarajy, Hartmut Witte

## Abstract

Accurate detection of tumors is critical for the success of oncologic surgical intervention. Along with any modality for the localization of tumors, surgeons achieve important information with direct palpation as long as direct touch of the tumor is possible. However, often due to using surgical tools in confined cavities or employing robotic end-effectors, direct palpation is not possible. We propose a low-frequency vibro-acoustic vibration sensing method to identify tumors based on their mechanical properties. The tumor detection problem was cast into a binary classification as a complementary modality. The method involves a wavelet-based multilayer perceptron neural network that is trained in a supervised manner. Some phantoms of healthy tissue with tumor model inclusion were used for experiments and data collection. From the 120 overall number of experiments, 18 data (15% of the whole data samples) were used as test data to evaluate the performance. The results show an 83.3% accuracy related to the confusion matrix, reflecting the model performance on previously unseen data. Performance of the classifier was evaluated using the confusion matrix and the receiver operating characteristic (ROC). It is discussed that the advantage of the proposed system is that the sensor is not directly touching the specimen and can be integrated into the probe end due to its small dimension.

## 1. Introduction

Identification and localization of abnormal cell growths, particularly cancerous tumors, within the human body is referred to as tumor detection. Quick and correct tumor detection significantly improves treatment outcomes, enhances survival rates, and minimizes further invasive procedures. Advances in imaging and diagnostic technologies have allowed for earlier interventions, reducing the likelihood of metastasis and improving patient prognoses. In particular, intraoperative tumor detection is a critical aspect of cancer surgery, ensuring complete tumor removal while preserving as much healthy tissue as possible. Real-time detection during surgery aids in reducing recurrence rates and improving functional outcomes. Surgeons rely on various imaging and sensing techniques, including fluorescence imaging, Raman spectroscopy, and ultrasound (US), to achieve precise tumor delineation. This real-time feedback is particularly important in neurosurgery, breast cancer surgery, and other procedures where tissue conservation is vital.

Several methods have been proposed for tumor detection, each with its advantages and limitations. Imaging-based methods such as magnetic resonance imaging (MRI) [1] and computed tomography (CT) [2] are widely used for this aim because of their detailed anatomical information output. MRI offers high-resolution images with excellent soft-tissue contrast, making it suitable for brain and soft-tissue tumors, but it is expensive and time-consuming. On the other hand, CT scan systems can deliver rapid imaging, though they involve exposure to ionizing radiation. Positron emission tomography (PET) complements these techniques by offering functional imaging based on metabolic activity, which is particularly useful for detecting highly dynamic cancerous cells, with the limitation of lower spatial resolution. US imaging is another modality [3] that provides real-time imaging without radiation exposure, but its usefulness can be limited by operator dependency and resolution constraints.

In addition to imaging methods, optical and molecular techniques have been explored for tumor detection applications. Fluorescence-guided surgery (FGS) [4] makes use of contrast agents that selectively bind to tumor cells and enables surgeons to visualize cancerous tissues intraoperatively. While this technique enhances surgical precision, it requires implementing exogenous dyes. Raman spectroscopy, another molecular-level technique, provides specific biochemical information, although it is complex and requires sophisticated equipment.

As cancerous cells possess higher cellular proliferation, altered extracellular matrix composition, and abnormal vascularization, the tumors exhibit altered physical properties compared to their normal tissues. Such physical properties include mechanical (e.g. elasticity, density, and viscosity), optical, and electrical (e.g. conductivity or impedance) properties. Tissues can become stiffer due to increased collagen deposition and crosslinking, but as well more compliant due to central necrosis provoked by malnutrition. Differences in light absorption and scattering enable optical imaging techniques to distinguish tumors from normal tissue [5]. Moreover, cancerous cells often have altered ion concentrations, affecting their electrical properties, which can be utilized in impedance-based detection [6] [7].

The mechanical properties may play the role of a basis for detection methods like elastography and mechanical palpation [8,9]. These techniques exploit the fact that the stiffness of cancerous tissues is meaningfully different from that of the healthy counterparts. For example, in elastography, US or MRI is employed to measure tissue displacement under mechanical stress, and the stiffness maps can be displayed quantitatively [10]. Tactile systems can assist surgeons with tumor localization through force-feedback mechanisms [11]. When surgeons have to use mechanical tools instead of directly touching of the patient’s organs, a mechatronic sensor or transducer may help to reflect the palpation and some mechanical properties of the tissue.

Nevertheless, measurements from biological systems yield complex time signals, and the raw data obtained directly from sensors exhibit non-stationary and unstructured behavior. For this reason, spectral analysis techniques are widely employed in biomedical imaging and diagnostics to extract features associated with pathological changes, enhancing the sensitivity and specificity of detection tools. Particularly, time-frequency analysis plays a significant role in characterizing complex biological signals such as those acquired during tumor detection. This approach analyzes how the frequency content of a signal evolves, making it principally suitable for non-stationary signals. Standard methods include the short-time Fourier transform and the wavelet transform, offering multi-resolution analysis of varying frequency signals. Spectral analysis results, in the form of time-frequency images, offer a rich and structured representation of signal characteristics suitable for machine learning classification methods [12].

When transformed into image formats using techniques such as continuous wavelet transform (CWT), these spectral representations preserve localized features across scales and frequencies, which are particularly valuable for distinguishing hidden structures in non-stationary signals [13–15]. Perceptron-based neural networks (NN), including multilayer perceptron (MLP), are well-suited for classifying such data due to their capacity to learn nonlinear decision boundaries from input features [16]. By treating spectral images as high-dimensional feature vectors, MLP can leverage the intrinsic patterns within the transformed domain, enabling effective classification even with relatively simple network architectures. This approach is especially beneficial in applications involving vibration analysis, biomedical signals, or acoustic monitoring, where temporal dynamics play a crucial role in class differentiation.

In this work, we investigate a method for the detection of inclusions reflecting potentially malignant tumors within the tissue phantoms. The method of vibro-acoustic sensing followed by CWT and MLP classification is studied experimentally. This article is arranged as follows: The material and method are presented in Section 2. First, the motivation and necessity of the research are explained based on the authors’ intraoperative observations, leading to further investigations. Then, the proposed hardware and experimental setup for the empirical evaluation of the proposed method, followed by the underlying theory, are explained. The results are provided in Section 3, presenting the experimental results and the CWT MLP outcomes. In Section 4, the results are discussed by a comparison with US testing. Section 5 is devoted to the conclusion summarizing the research results.

## 2. Materials and Methods

This methodology was motivated by direct intraoperative observations conducted by the authors during surgical resections of head and neck malignancies. A surgeon employes monitors and an operating microscope, in conjunction with elongated surgical tools to navigate the restricted anatomical space and access the neoplasm. The mechanical impedance of the tumor is perceptible through tactile assessment, highlighting the potential utility of providing real-time haptic feedback to the operating surgeon. This observation served as a key motivation for developing a method to enable intraoperative tactile feedback.

Preserving excised tissue specimens for basic research purposes poses significant challenges, as biological tissues are prone to rapid degradation and cannot be stored for extended periods. However, when the primary focus is on mechanical characterization, synthetic elastomeric materials can be employed to replicate the mechanical properties of the resected tissues. In this study, some elastomeric tissue-mimicking phantoms were fabricated based on consultation with our medical experts to experimentally implement the proposed methodology. The phantoms were designed to replicate the mechanical contrast between healthy and tumor tissues.

The bulk of the phantom, representing healthy soft tissue, was constructed using Ecoflex™ silicone elastomer with a Shore hardness of 20A. To replicate a tumor inclusion, a stiffer material—Smooth-On™ Silicone 230 with a Shore hardness of A30—was embedded within the phantom. The elastic modulus of the Ecoflex-based healthy tissue analog was measured to range between 0.05 MPa and 0.10 MPa, whereas the tumor-mimicking inclusion exhibited a significantly higher modulus, ranging from 1.14 MPa to 1.49 MPa. This substantial stiffness contrast ensures a realistic mechanical differentiation between pathological and non-pathological regions, suitable for evaluating elastography imaging or tactile sensing systems.

Note that the Ecoflex™ silicone elastomer’s modulus range corresponds well to healthy tissues like fat (0.01-0.1 MPa), normal breast (0.02-0.06 MPa), liver (0.02-0.08 MPa), white matter of the brain (0.01-0.04 MPa), and muscle (at rest 0.02-0.05 MPa). On the other hand, the tumor inclusion analog (1.14–1.49 MPa) aligns with malignant breast tumors (e.g., invasive ductal carcinoma) (0.5–1.5 MPa), liver tumors (e.g., hepatocellular carcinoma) (1–2 MPa), prostate tumors (0.8–2 MPa), and fibrotic or calcified tissues (can exceed 1 MPa). The proposed vibro-acoustic method is schematically represented in Figure 1. Two microcontrollers and a PC control the hardware to collect the raw signal and perform further signal processing.

**Figure 1.**
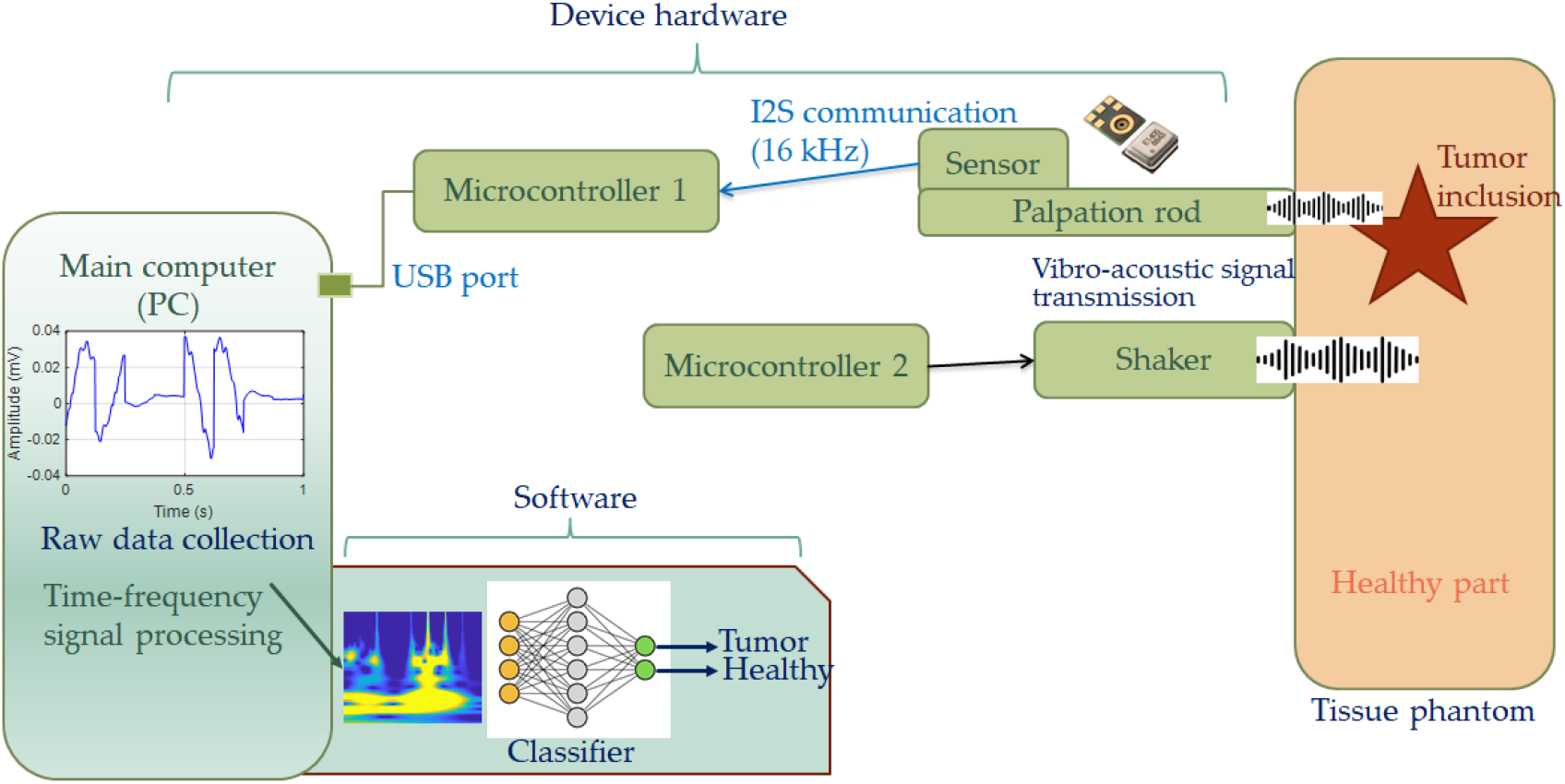
Schematic of the hardware and the proposed method.

Microcontroller 2 is used to generate vibro-acoustic vibrations as external excitations. The phantom is palpated through a 17 cm rod, representing a surgical tool or suction tube. At the end of the palpation rod, the implemented sensor transmits the response to the excitation to microcontroller 1 using a digital I^2^S protocol. The sensor is SPH0645LM4H-1-8, a digital micro-electro-mechanical systems (MEMS) microphone proposed in [17,18]. The main computer is a PC that collects the raw signal using a serial communication (with a USB port). The raw signal is processed using MATLAB^®^ R2022b (MathWorks^®^, Natick, MA, USA) using wavelet time-frequency analysis. In this study, we employ the CWT to accomplish time-frequency analysis of experimental signals collected from tissue-mimicking phantoms. The signals contain non-overlapping segments of 512 samples analyzed using a wavelet-based filter bank. The CWT provides a localized spectral decomposition of non-stationary signals, allowing simultaneous representation of time and frequency characteristics. Mathematically, the CWT of a signal *x*(*t*) is expressed as:

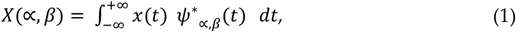

with the scaled and translated wavelet function defined by:

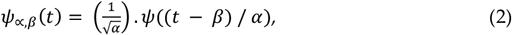

Where *α* > 0 is the scale parameter, *β* can be considered as the translation in time, and the function *ψ*(*t*) is the mother wavelet here adopted to be the analytic Morlet wavelet due to its optimal time-frequency localization [19]. In Matlab^®^ R2022b, the implementation was carried out using the *cwtfilterbank* object. The parameters were set to span a frequency range of 1–220 Hz, with a sampling rate 512 Hz and 12 voices per octave to ensure high frequency resolution. The output of the CWT consists of complex coefficients *X*(∝, *β*), from which we computed the magnitude scalogram:

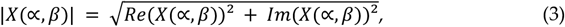

These scalograms were normalized via min-max normalization to standardize the data across segments:

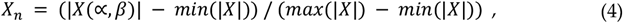

The resulting time–frequency images visualize spectral energy evolution and are suitable for input into machine learning pipelines, such as NN, for tissue classification tasks. A feedforward NN multi-layer perceptron, MLP, is employed for the classification task using MATLAB^®^ R2022b Neural Network Toolbox to distinguish between two classes. The MLP is created with a single hidden layer of 32 neurons, using the *patternnet* function specifically tailored for pattern recognition and classification problems. The training function selected is ‘*trainscg*’, scaled conjugate gradient (SCG), an efficient secondorder optimization algorithm that converges faster than basic gradient descent, especially for smaller datasets. The performance metric is set to ‘*crossentropy*’, which is ideal for classification tasks because it measures the divergence between the predicted and actual class probabilities.

The MLP takes a feature vector, *X* ∈ ℝ^*n*^, as input, processes it through a hidden layer with the log-sigmoid activation function, and then the class probabilities at the output layer are estimated using its *softmax* function. We take *W*_*1*_ and *b*_*1*_ vectors as the weights and biases of the hidden layer, and *W*_*2*_, *b*_*2*_ for the output layer. Then, our forward propagation rule is described by the following equations:

Hidden layer can be expressed as

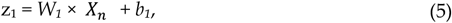

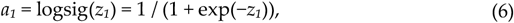

Likewise, the output layer (*softmax* for two-class classification) is given as

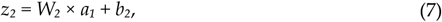

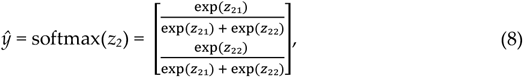

Within the network training procedure, the cross-entropy loss, given below, is targeted for minimization

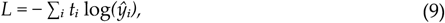

where *t*_*i*_ is the one-hot encoded true label and *ŷ*_*i*_ is the predicted probability for class *i*. The parameters *W*_*1*_, *W*_*2*_, *b*_*1*_, and *b*_*2*_ are optimized using the SCG algorithm to minimize this loss over the training dataset. The dataset is randomly divided into three subsets using *net*.*divideFcn = ‘dividerand’*, where 75% of the data is allocated for training (*trainRatio* = 0.75), 10% for validation (*valRatio* = 0.1), and 15% for testing (*testRatio* = 0.15). This random partitioning helps to ensure the network is trained on a representative portion of the data, validated to monitor overfitting during training, and tested to evaluate the model’s ability to generalize to unseen samples. This structured approach allows the network to learn effectively while providing insight into both its learning behavior and real-world predictive performance.

## 3. Results

Figure 2 shows the proposed experimental setup. The system contains one Nodemcu ESP 32 microcontroller to generate arbitrary excitations employing a PAM8302A amplifier and a Feather M0 microcontroller to transmit the sensor data to the PC using a USB port. The palpation rod is a 4 mm diameter solid aluminum bar placed freely over the phantom.

**Figure 2.**
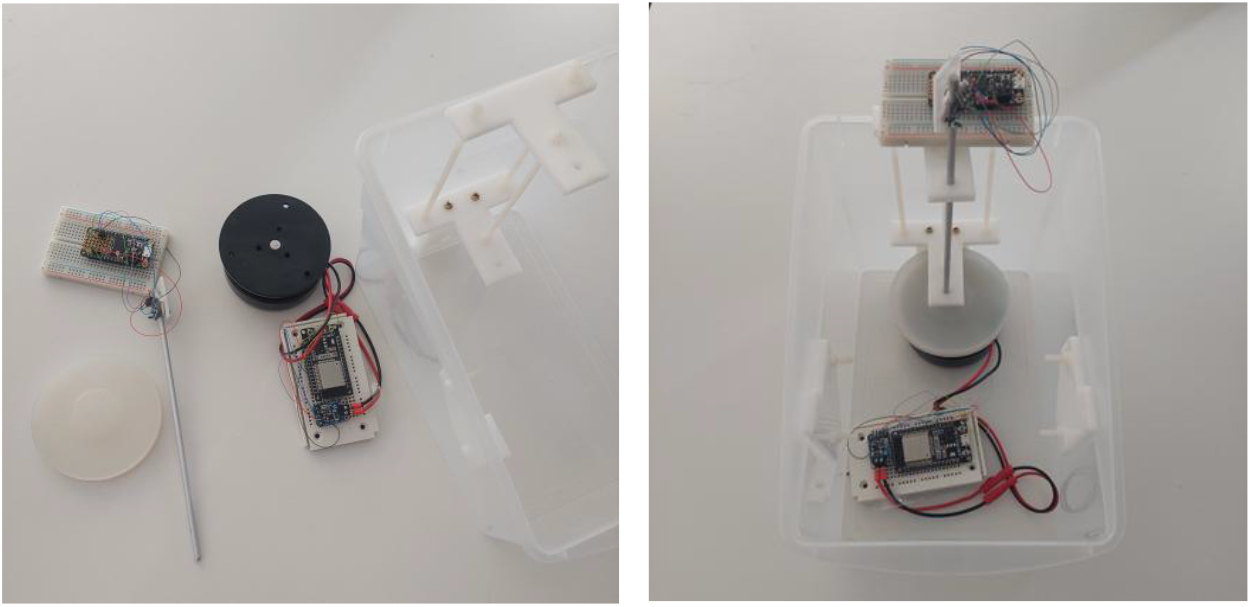
The experimental setup.

In each experiment, involving a 1s data collection, the shaker produces an excitation of frequency 130-*n* Hz for 500ms, where 1<*n*<10 is the number of experiments with a sample, and then a tone of frequency 110+*n* Hz for another 500ms. Note that with 6 ‘healthy’ and 6 ‘tumorous’ samples, the total number of experiments is (6+6)*10=120. The system time response was collected by the Feather M0 microcontroller and then sent to the PC for each experiment. Ten typical recorded time signals are shown in Figure 3 for both tumor and healthy cases. For visual comparison, the data obtained from testing tumorous and healthy phantoms are given in common plots. Nevertheless, differentiation of the time signal waveforms does not seem to be trivial at all. The graphs represent unsteady behavior that makes the signal analysis and comparison difficult.

**Figure 3.**
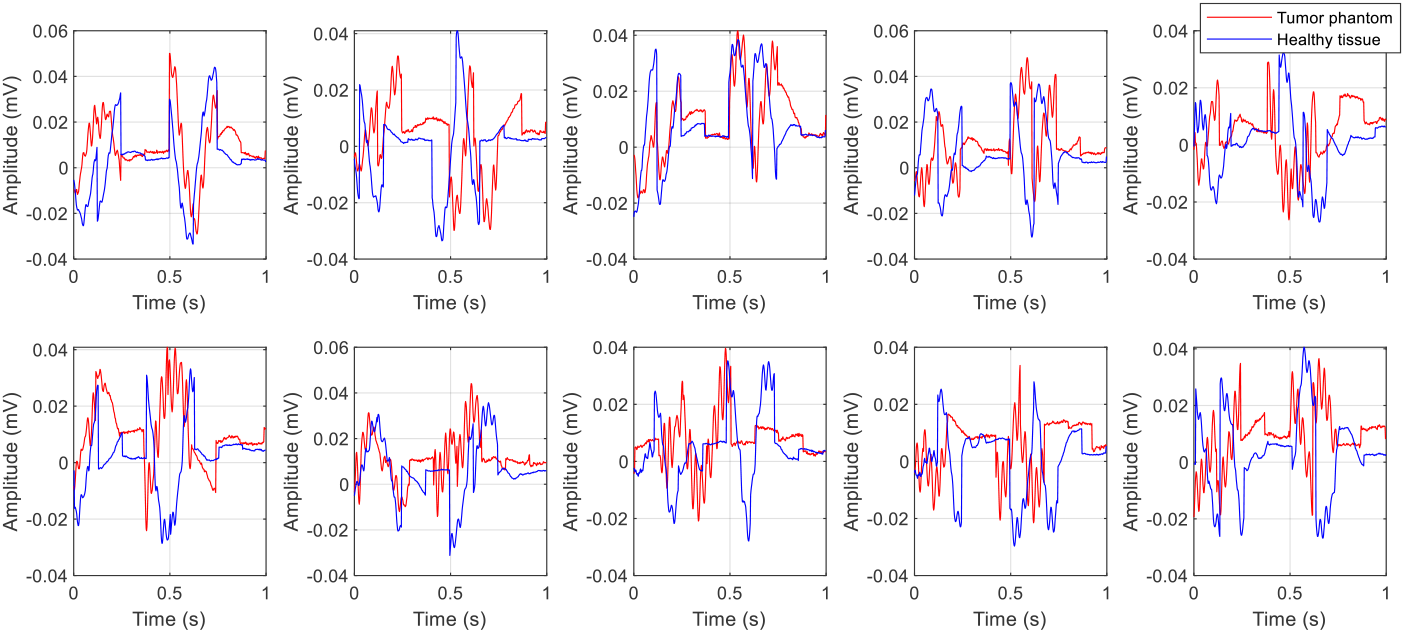
Time history of experimental data collected from tumorous and healthy tissue samples (the plots correspond to only 10 typical experiments).

Therefore, a CWT analysis is employed to transform the raw data into a format that is more suitable for either visualization or machine learning applications. The results shown in Figure 4 reveal more structured images that can be further analyzed with NN classification methods, MLP in this case. The classification results indicate that the trained MLP achieved an overall accuracy of 85.8%, meaning that across the entire dataset, including training, validation, and test sets, the model correctly classified 85.8% of the samples. However, the test accuracy is slightly lower at 83.3%, which reflects the model’s performance on previously unseen data. This small gap between overall and test accuracy suggests that while the model has learned the training data well, its generalization to new, unseen samples is slightly limited. The model performs reasonably well, but improvements could be achieved by increasing the training data, tuning the network architecture (e.g., regularization or dropout), or experimenting with different feature extraction methods.

**Figure 4.**
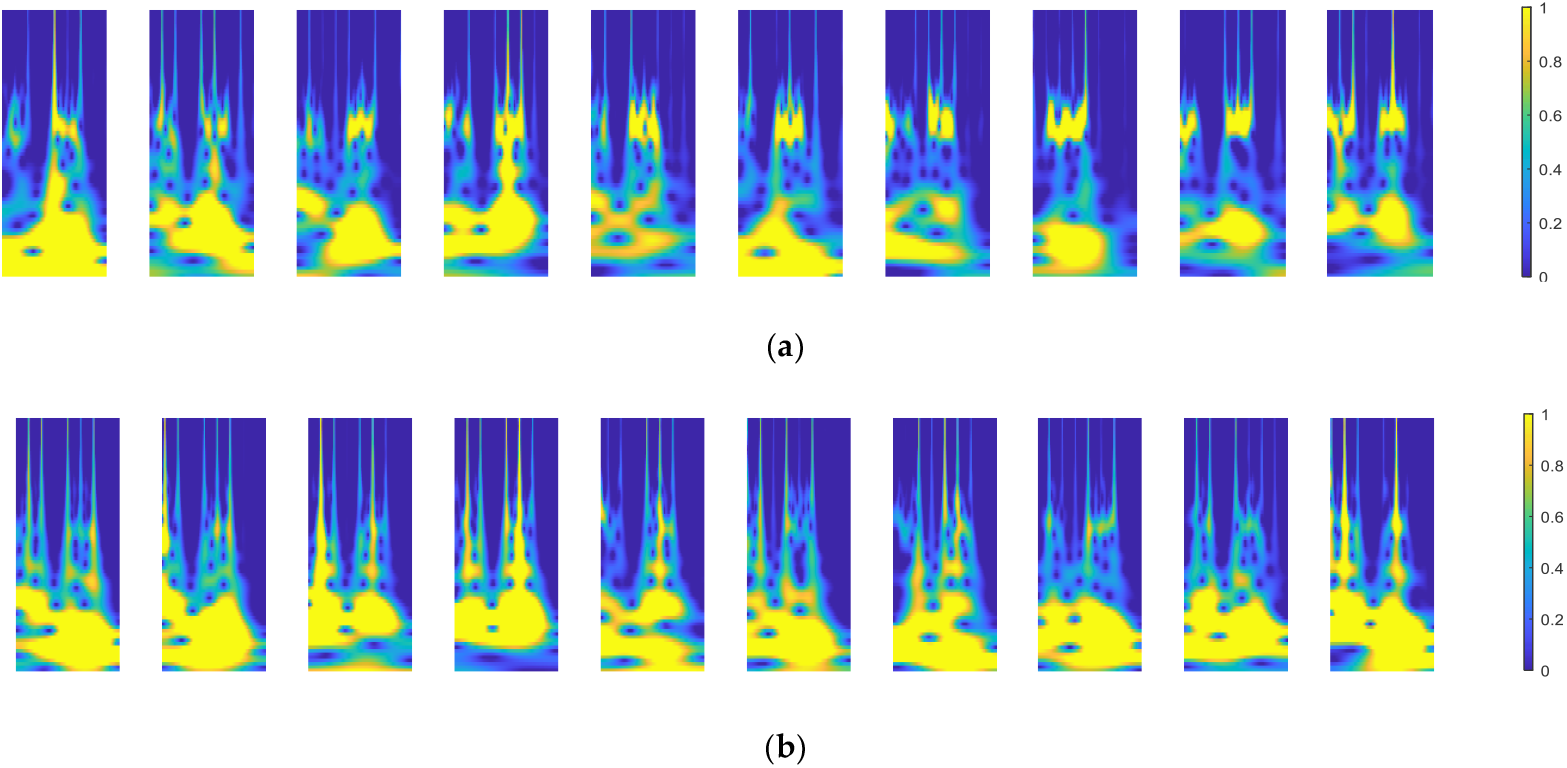
The CWT results for the typical experiments: (**a**) Tumor phantom data; (**b**) Healthy tissue.

Nonetheless, the results show that the classifier is capable of distinguishing between the two classes with good reliability. Performance of the classifier is analyzed using the confusion matrix and the receiver operating characteristic (ROC). The confusion matrix is a visual summary of the classifier’s performance on the test set, distinguishing between two classes: Class 1 (Tumor) and Class 2 (Healthy). The matrix given in Figure 5a shows that out of the 9 actual Tumor samples, 7 were correctly predicted as Tumor and 2 were misclassified as Healthy, giving a true positive rate of 77.8% for Tumor detection. Additionally, out of 9 actual Healthy samples, 8 were correctly classified, while 1 was misclassified as Tumor, resulting in an accuracy of 88.9% for identifying Healthy cases.

**Figure 5.**
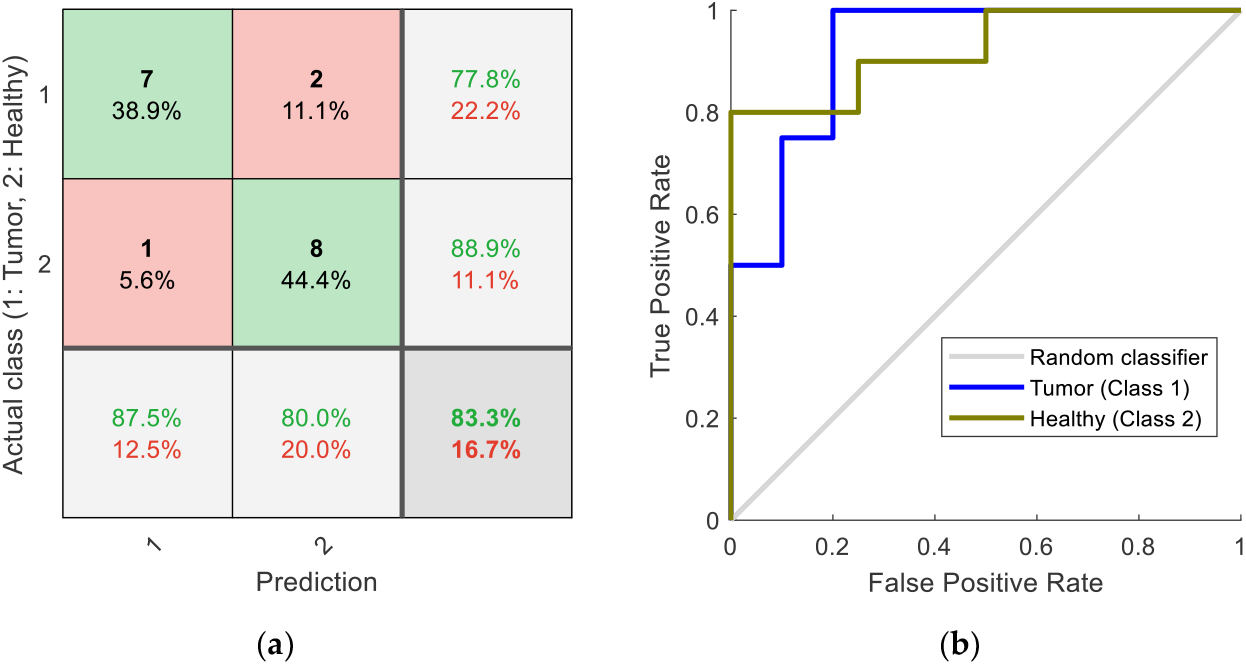
Performance of the classifier with test data: (**a**) The confusion matrix; (**b**) The ROC curve.

The ROC curve is a graphical tool used to assess the performance of a binary or multiclass classifier by plotting the true positive rate (TPR) against the false positive rate (FPR) at various classification thresholds. A good classifier achieves a TPR close to 1 while maintaining a low FPR, thus pushing the ROC curve toward the top-left corner of the plot.

The diagonal gray line represents the performance of a random classifier, which is the baseline for comparison. In the ROC plot given in Figure 5b, two curves are given for Class 1 (blue) and Class 2 (olive green). Both curves show a strong upward trajectory, indicating good classification performance. The blue curve for Class 1 reaches the maximum TPR of 1.0 at a low FPR, suggesting that most positive samples are correctly classified with few false alarms. Similarly, the curve for Class 2 also demonstrates strong performance.

## 4. Discussion

To discuss the method and the results from the perspective of current commercial technology, it would be interesting to examine how the inclusions within elastomeric phantoms can be demonstrated using US-based methods, similar to those performed in non-destructive testing (NDT) techniques [20]. One such system involves the Vantage 128 from Verasonics, which is equipped with an array of US transmitters and receivers. In a typical setup, the transmitter emits focused ultrasonic pulses into the phantom material. As the wave propagates, it interacts with various internal features, including boundary interfaces and embedded inclusions. These interactions cause partial reflections of the wave, which are subsequently captured by the receiving elements of the array. By analyzing the time-of-flight and amplitude of the received signals, the system can reconstruct an image or signal profile that indicates the presence, location, and acoustic contrast of the inclusion in soft-tissue mimicking materials.

The Vantage 128 from Verasonics Inc. is a research-grade US data acquisition system featuring 128 transmit and receive channels, widely used in NDT and medical imaging research. The test shown in Figure 6 reveals the inclusion within the phantom. However, such systems are bulky and need many parameter adjustments as they are used for research on a wide range of materials. A new and portable device specifically designed for biomedical applications is the handheld US imaging system, iQ3 device, from Butterfly Network Inc. Powered by semiconductor-based Ultrasound-on-Chip™ technology, the iQ3 offers high-resolution imaging with a single probe and can be tethered to smartphones or tablets. The experiment with an iQ3 device is shown in Figure 7. The inclusion or the tumor model is observed as a white line on the screen. Nevertheless, the US systems still have bulky probes and need to touch the specimen surfaces.

**Figure 6.**
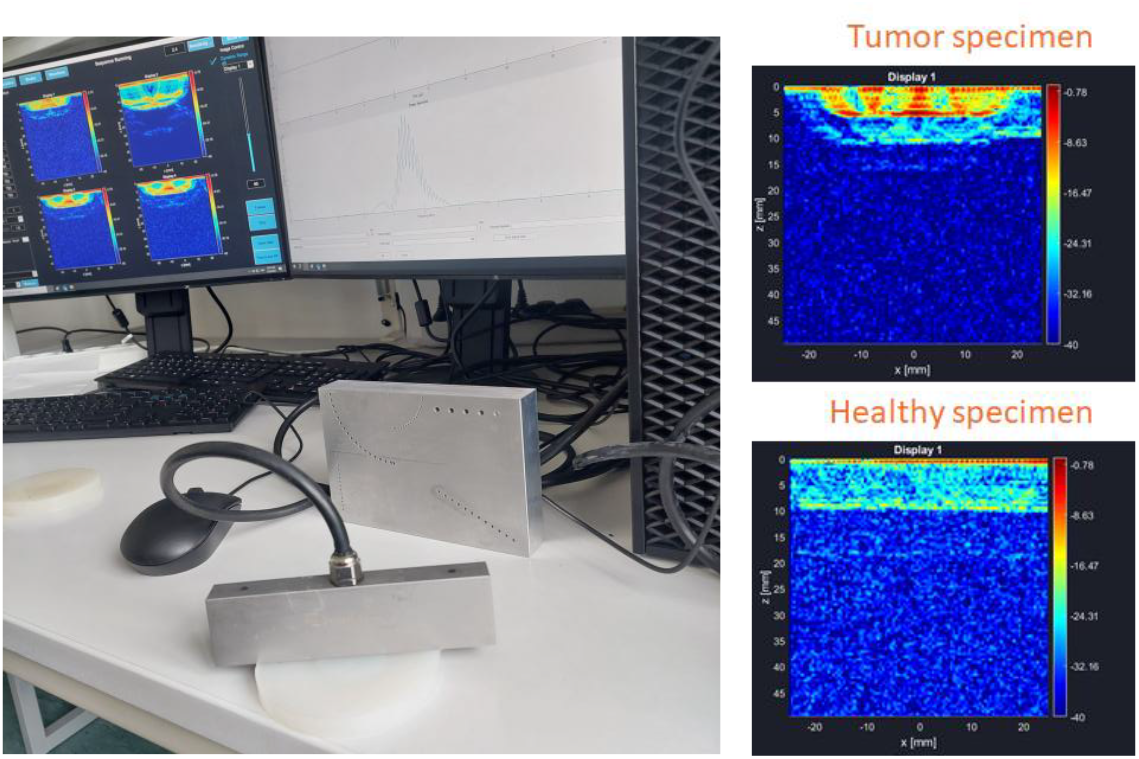
Typical US examination of the phantoms.

**Figure 7.**
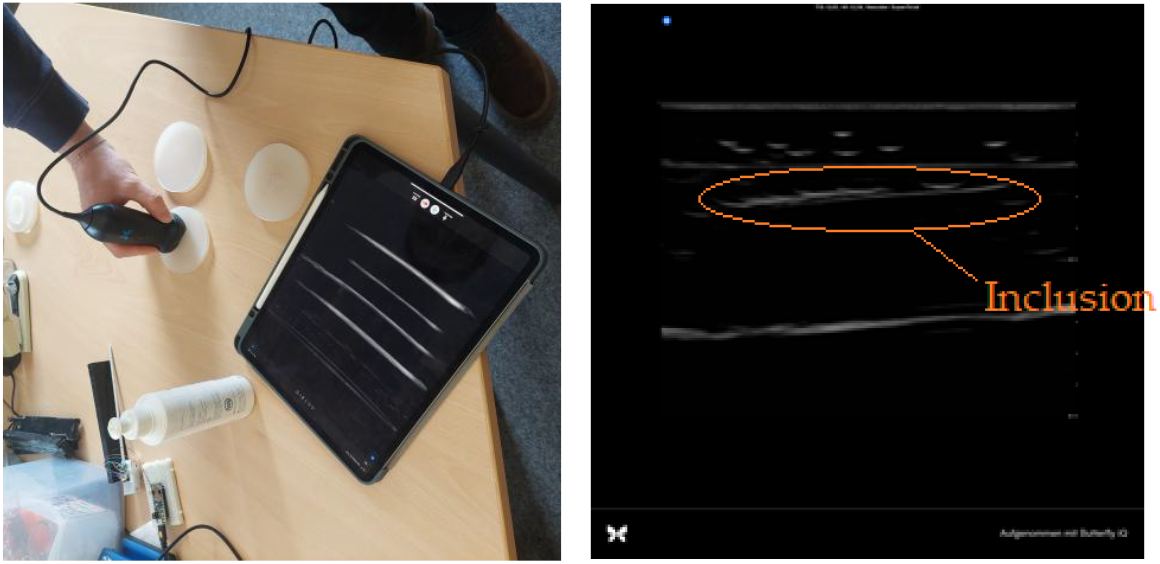
The US examination using iQ3.

In medical applications where surgical tools (such as a suction tube or manual or robotic grippers) are operated on the patient’s body, a small sensor indirectly performing pointwise measurements is required. In contrast, the vibro-acoustic method is based on relatively low-frequency forced vibrations, sensing indirectly at the end of a tool (a rod in this work). The system works with data packages collected in 1s periods. While the time signals do not represent comprehensible gesticulations, the CWT provides images showing more structured profiles that are efficiently recognizable by the MLP classifier. Totally 120 wavelet images (60 ‘tumor’ and 60 ‘healthy’) were recorded and labeled for offline training of a feed-forward NN, the MLP classifier. The confusion matrix of the trained MLP network indicates that the overall classification accuracy is high, 83% as seen in the bottom-right cell of Figure 5a, and the matrix indicates that the model is more accurate in identifying healthy cases than tumor cases. The false negative rate (‘Tumor’ predicted as ‘Healthy’) is slightly higher than the false positive rate, which may be important if the cost of missing a tumor is high.

The color coding in the matrix helps highlight performance: green for correct predictions and red for misclassifications, giving an intuitive view of classifier reliability. Additionally, since both ROC curves, in Figure 5b, are well above the diagonal line, we can conclude that the classifier distinguishes between the two classes efficiently. From the NN point of view, the results confirm that the MLP classifier is performing well with good sensitivity and specificity.

## 5. Conclusions

In this article, the need for a measurement system reflecting the mechanical properties of a tumor in cancer tumor removal was introduced. The newly introduced method based on vibro-acoustic measurement was proposed to develop a system that can be integrated into thin surgical tools or probes for tumor detection. The feasibility of the proposed method was examined experimentally using artificial phantoms. A wavelet-based neural network method, CWT MLP, was proposed to classify the collected data into healthy or tumor classes. The analysis shows promising results in the feasibility of the detection of tumors, which can help the surgeon as a supplementary modality. It was discussed that in comparison with US systems, the proposed method has the advantage of small sensor size and integrability with tools (indirect touch of the specimen). Improvement of the hardware, e.g. by employing more sophisticated microcontrollers, integrating the sensor with actual surgical tools, and in particular, experiments with real tissues and tumors for collecting data and retraining the method, remains a proposal for future work.

## Author Contributions

Conceptualization, M.S. and H.W.; methodology, M.S.; software, M.S.; validation, M.S. and H.W.; formal analysis, M.S.; investigation, M.S. and H.W.; resources, H.W.; data curation, M.S.; writing—original draft preparation, M.S.; writing—review and editing, M.S. and H.W.; visualization, M.S.; supervision, H.W.; project administration, H.W.; funding acquisition, H.W. All authors have read and agreed to the published version of the manuscript.

## Informed Consent Statement

The study did not involve humans.

## Acknowledgments

The authors would like to express their gratitude to Prof. Florian Römer from Fraunhofer Institute for Non-Destructive Testing IZFP for technical support in the US measurement.

## Abbreviations

The following abbreviations are used in this manuscript:

CT: Computed tomography
CWT: Continuous wavelet transform
MEMS: Micro-electro-mechanical systems
MLP: Multilayer perceptron
MRI: Magnetic resonance imaging
NDT: Non-destructive testing
NN: Neural networks
PET: Positron emission tomography
ROC: Receiver operating characteristic
SCG: Scaled conjugate gradient
TPR: True positive rate
US: Ultrasound

